# Impacts of diets fed after weaning on gut microbiota and susceptibility to DSS-induced colitis in mice

**DOI:** 10.1101/549501

**Authors:** Yu Meng, Xiaojun Li, Shuijiao Chen, Fujun Li, Yani Yin, Jianping Liu, Fanggen Lu, Xiaowei Liu

## Abstract

**Background:** Living in a sanitary environment and taking Western-style diet in early life are both risk factors for inflammatory bowel disease and important factors for shaping host gut microbiota. Here, we aimed to establish whether different dietary composition fed during the early period after weaning would associate the susceptibility to DSS-induced colitis with different gut microbiota shifts.

**Methods:** Eighty weaned Balb/c mice were fed with high sugar, fat, protein, fiber, and standard diet for 8weeks respectively. Inflammation was induced by administration of 2.5% (wt/vol) dextran sulfate sodium (DSS) in drinking water for 7 days, and the microbiota characterized by 16s rRNA based pyrosequencing. Analyzed the inflammatory factors and toll-like receptors by Real-time PCR

**Results:** The high protein and high fiber+protein group exacerbated severity of DSS-induced colitis, the high fiber and high protein+fiber groups had the effect of reducing colitis, and the high sugar, fat and standard group show the similar disease phenotype of colitis. The diversity and richness of the microflora were significantly decreased in the high fiber group, while only decreased richness of flora was observed in the high protein group. The abundance of *Firmicutes* was decreased and the abundance of *Bacteroides* was increased in the high fat, high sugar, high protein and high fiber groups, especially in the high protein and high fiber group. The microbial community structure was slightly different at the species/genus level. The microbial community structure of high protein-fiber group and high fiber-protein group was still similar.

**Conclusions:** Mice were fed with different dietary compositions of high sugar, fat, protein and fiber diets since weaning, and similar gut microbiota of high-abundance *Bacteroides* and low-abundance *Firmicutes* are formed in adult mice. These microbiota do not cause colonic mucosal damage directly. Only high protein diet aggravated DSS-induced colitis, while high fiber diet alleviated the colitis.

## Introduction

The pathogenesis of inflammatory bowel disease (IBD) is considered to be intolerance of the genetically susceptible individuals to gut microbiota, triggering a dysregulated immune response resulting in inflammatory injuries of intestinal mucosa ^[1–3]^. Epidemiologic data show that the incidence of IBD among first-generation immigrants to regions with high incidence of IBD is as low as that among natives in the countries from which they emigrated, yet second-generation immigrants have an increased risk of IBD similar to that in locations to which they immigrated ^[4]^. The variance in the disease incidence cannot be solely attributed to genetic susceptibility, suggesting environmental factors exposed in early life play important roles in the development of IBD ^[4,5]^. Current evidences support risk factors associated with IBD include hygiene hypothesis, Cesarean section, formula feeding, and antibiotic use in infants and young children ^[6–8]^. Differences in delivery mode, infant feeding type, sanitary condition and antibiotic use have also been linked with differences in the intestinal flora of babies and children ^[9]^. Several studies have demonstrated marked alterations in the gut microbiota of patients with IBD ^[10–12]^. Thus, the composition of gut microbiota is associated with IBD.

Studies have found significant differences in fecal microflora between people eating western diet and people from regions where fiber content is high in the diet, revealing impacts of different diets in shaping gut microbiota ^[13,14]^. Recent reports indicate that the gut microbiota and alterations thereof, due to a consumption of a diet high in saturated fats and low in fibers, can trigger factors regulating the development of IBD ^[15]^. The incidence of IBD is steadily rising in developed as well as in developing countries paralleling the escalating consumption of western-style diets, characterized by high protein and fat as well as excessive sugar intake, with less vegetables and fiber ^[16]^. Animal studies also find that “western-style” diets can lead to intestinal flora shifts in mice and aggravate inflammatory injuries of colonic mucosa induced by DSS ^[17]^. All these findings suggest that changes in dietary composition contribute to the development of IBD by altering the structure of gut microbiota.

In the recent years, the incidence of pediatric IBD is rapidly on the rise, indicating that gut microbiota shaped in early life is related to the risk of IBD ^[18]^. The development of intestinal flora consists of two stages, one is lactation and the other is exposure to environment and diet after weaning ^[19]^. A comparative study has found similarities in fecal microflora between European children of preweaning and children of the same age from rural Africa, and significant differences between European children aged 2 to 6 years and children of same age from Africa ^[20]^. A recent analysis of the intestinal flora composition of one infant followed over 2.5 years showed a considerable alteration in the microflora with the introduction of solid foods and a shift towards a more stable, adult-like microbiota with weaning ^[21]^. By 3 years of age, microbiota composition approaches that of adults ^[21]^. These results indicate that the development of intestinal flora after weaning is influenced by diets. Thus, the development of gut microbiota during the early period after weaning may be related to the development of IBD.

Taken together, we propose a hypothesis that different dietary composition fed during the early period after weaning may affect the risk of the development of IBD by different gut microbiota shifts. Therefore, the aim of this study is to investigate the developments of gut microbiota compositions in mice fed on different diets after weaning as well as the susceptibility to DSS-induced colitis.

## Methods

### Animals, diets and experimental design

A total of eighty weaned female Balb/c mice (weight 12±0.8 g) were caged under standard conditions (20-22°C, 50-55% humidity, 12/12h dark/light cycle), and randomly assigned to eight groups (10 mice in each group): control group (Con), standard group (Sta), group with high sugar (Sug), group with high fat (Fat), group with high protein (Pro), group with high fiber (Fib), group with one-month high protein and one-month high fiber (Pro+Fib), group with one-month high fiber and one-month high protein (Fib+Pro) (Table 1). All mice were provided with water and diets, and the calories were balanced in all the groups every day. The stool of mice was collected once after 8-week treatment and stored at −70°C for further analysis. Susceptibility to colitis was tested by the administration of 2.5% (wt/vol) dextran sulfate sodium (DSS; Molecular Weight 36000-50000, MP Biomedicals, LLC, Solon, OH, USA) in the drinking water for 7 days after 8-week treatment. The animals were weighed weekly and the length of colon was measured following CO^2^ asphyxiation at the end of experiments. This study was approved and monitored by the Ethics Committee of the 2nd Xiangya Hospital of Central South University, China.

**Table 1.**
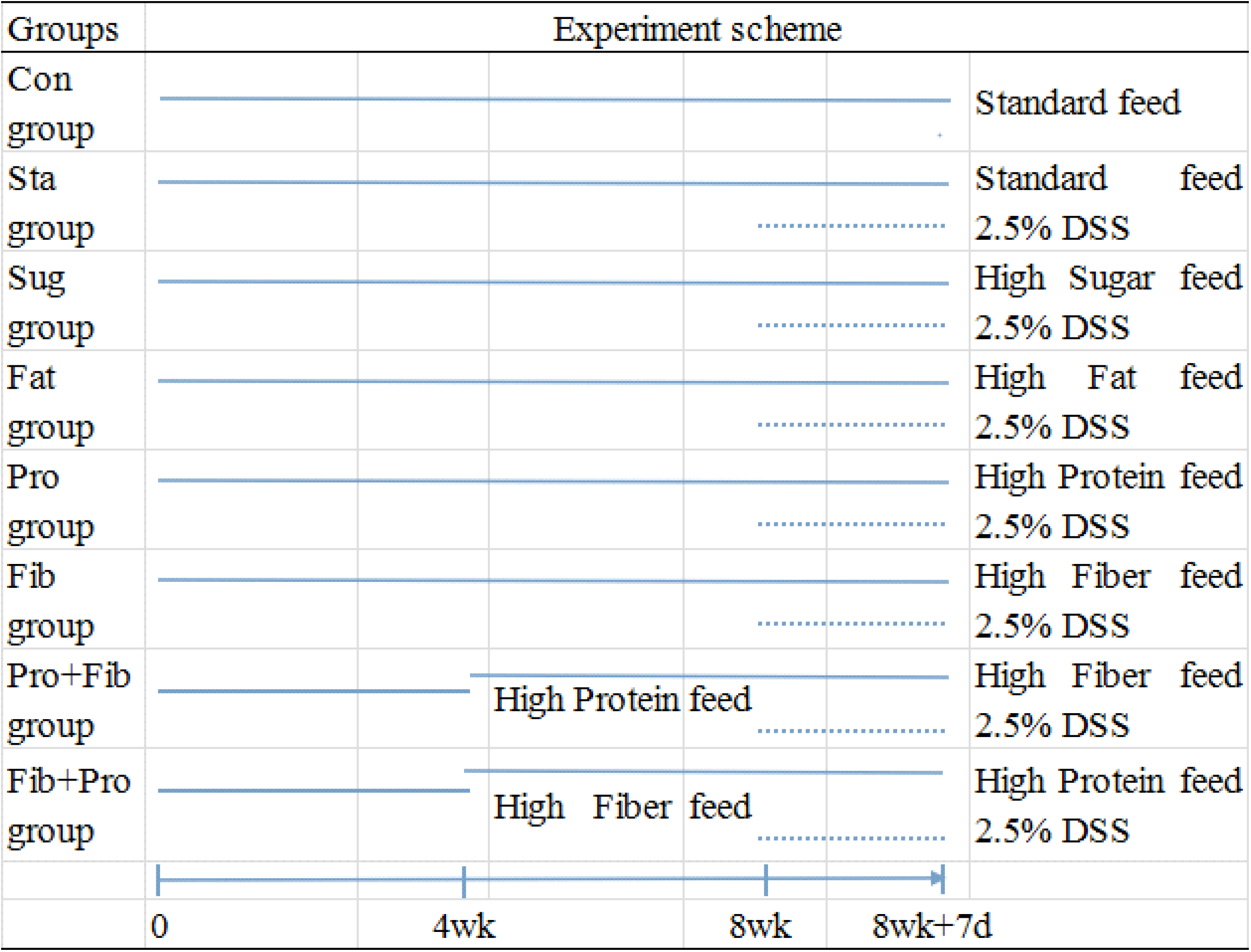
The experiment scheme of food intervention.

### Disease activity index

Disease activity index (DAI) was quantified using the parameters of weight loss, stool consistency, and gross blood in the feces which were recorded daily for each animal ^[22]^.

### Histological analysis

Colons were immediately excised, rinsed with ice-cold phosphate-buffered saline. 2 sections were snap-frozen in liquid nitrogen and stored at −70°C for RNA isolation and protein preparation. Tissue sections from the distal colon were fixed in 10% buffered formalin and embedded in paraffin. The sections were cut as 6 mm and stained with hematoxylin and eosin (H&E).

### RNA extraction and quantitative RT-PCR

Total RNA from colonic tissue samples was extracted using the EastepTM Total RNA Extraction Kit (Promega) according to the manufacturer’s protocol. RNA concentration was determined spectrophotometrically. The RNA was reverse transcribed into cDNA using the GoScriptTM Reverse Transcription System (Promega). Quantitative real-time polymerase chain reaction (PCR) was performed for CGRP with GoTaq^®^ qPCR Master Mix (Promega) using the DNA-binding dye, carboxy-X-rhodamine (CXR) reference dye (identical to ROX™ dye), for the detection of PCR products on a ViiATM7 Real-Time PCR system (ABI).

Amplification of MUC2, TLR2, TLR4, TLR5 and glyceraldehyde phosphate dehydrogenase (GAPDH) was routinely performed. The PCR amplification profiles consisted of hot-start activation at 95°C for 10 min, followed by cycles of denaturation at 95°C for 15 s, annealing at 60°C for 30 s and extension at 72°C for 15 s. A dissociation stage was added at 95°C for 15 s, 60°C for 15 s, and 95°C for 15 s. Melting curve analysis of amplification products was performed at the end of each PCR reaction. Each PCR reaction was performed in triplicate, and the mean Ct value was calculated. Relative fold changes in RNA levels were calculated by the △△Ct method.

### Pyrosequencing of fecal microbiota and bioinformatic analysis

Genomic DNA was extracted from the stool of enrolled mice (3 mice per group) using the QIAamp DNA Stool Mini kit (Qiagen, Germany). The extracted DNA was amplified using primers targeting the V1 to V3 hypervariable regions of bacterial 16S rRNA gene (27F: 5’-AGAGTTTGATCCTGGCTCAG-3’, 533R: 5’-TTACCGCGGCTGCTGGCAC-3’). PCR reaction mixtures were denatured at 95°C for 2 min followed by 25 cycles of denaturation at 95°C for 30 sec, annealed at 55°C for 30 sec, and elongated at 72°C for 30 sec. DNA sequencing was performed using a 454 GS FLX Titanium Sequencing System (Roche, USA).

Mothur software (Mothur v.1.25.1; http://www.mothur.org) was used to process the sequence data ^[23]^. Sequencing reads from the different samples were sorted by unique barcodes. Primers and barcodes were trimmed from each read. To minimize the sequencing errors, chimeric sequences were removed using UCHIME (http://drive5.com/uchime). Low-quality sequences were excluded if the sequences met one of the following criteria: (1) sequences with more than two inexact match to forward primer; (2) sequences with unrecognizable reverse primer; (3) sequences with a length of < 200 nucleotides; (4) sequences with undetermined bases in the sequence read; (5) sequences with average quality < 25. The trimmed sequences were clustered into operational taxonomic units (OTUs) by setting a 0.03 distance limit (97% of similarity). The alpha diversity (Shannon index and Simpson index) and richness (ACE and Chao1) were analyzed based on identified OTUs using the Mothur program. Representative sequence of each OTU was taxonomically classified using the Silva database^[24]^ at a 70% confidence threshold with Mothur.

## Results

### Effects of different diets on the expression of MUC2, TLR2, TLR4, TLR5, IL-1β and IL-6

The expression of MUC2, TLR4 and TLR5 had no difference in all groups before adding the DSS (Fig 3, P > 0.05). The expression of TLR2 in the Sug group was higher than the other groups before adding the DSS (Fig 3D, P < 0.05), had no difference between the Pro group and the Fib group (Fig 3D, P > 0.05), and had no difference between the Pro+Fib group and the Fib+Pro group (Fig 3D, P > 0.05). After adding the DSS, the expression of IL-1β was highest in the Pro group, and had significant difference with the other groups (Fig 3A, P < 0.05). The expression of IL-6 was lowest in the Fib group, and had significant difference with the other groups (Fig 3B, P < 0.05).

### Effects of different diets on DSS induced colitis in the mice

After 7 days of DSS administration, the mice in the Sta, Sug, Fat, Pro and Fib+Pro groups developed significant weight loss, diarrhea, rectal bleeding, and physical weakness compared to the control mice treated with PBS. The mice in Pro group and Fib+Pro group showed significantly aggravated colitis compared with the other groups (P < 0.05), with more colon shrinking, more body weight loss and higher DAI. However, little DSS-induced bleeding and significantly less colon shrinking or DAI were observed in the Fib or Pro+Fib groups (Fig 1, 2). DSS induction resulted in the epithelial destruction, edema, ulcerations, goblet cell depletion, and intense inflammatory infiltration compared to non-DSS-treated controls, with the most serious effects in Pro group.

**Fig 1.**
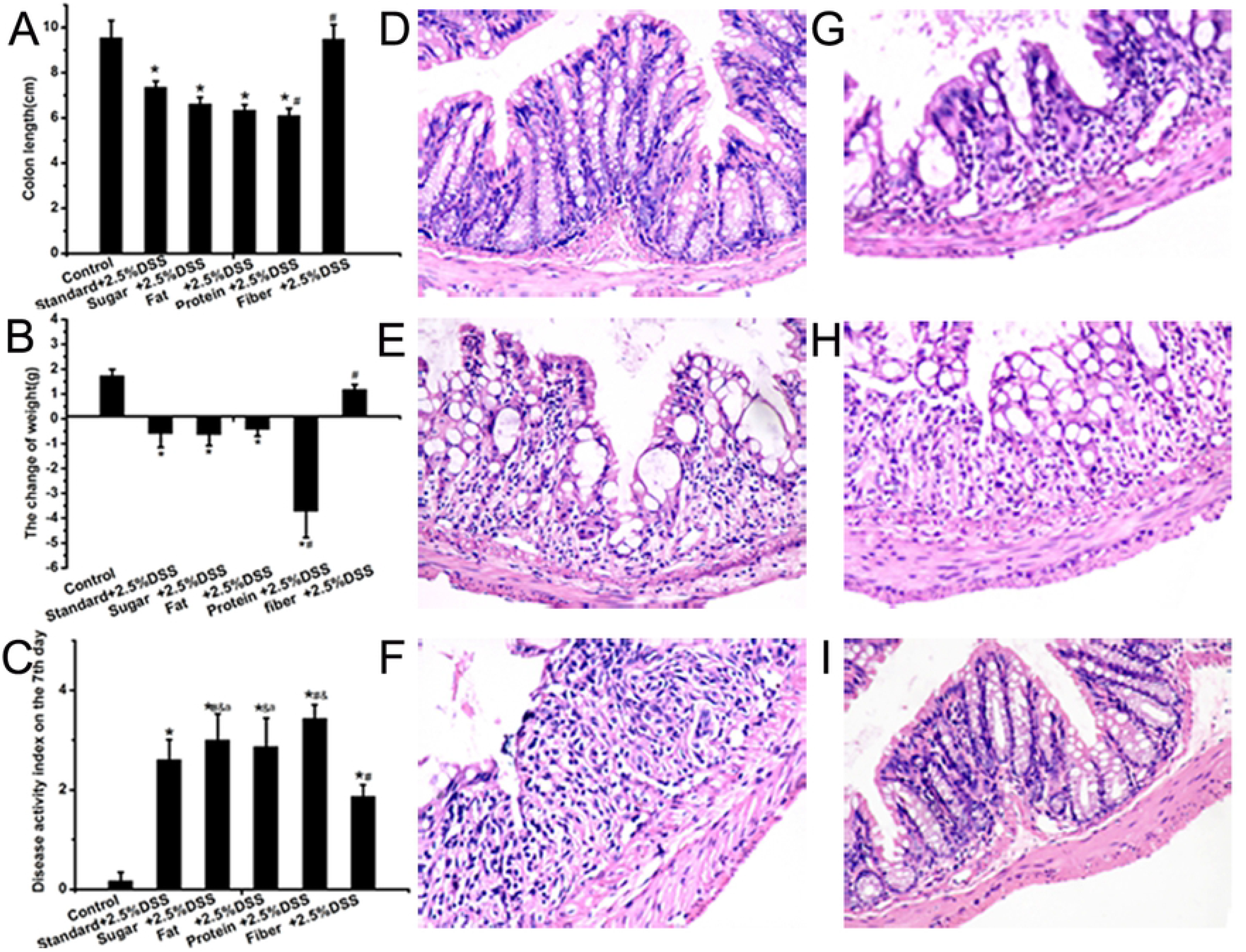
Effects of different diets on DSS induced colitis in the mice. Five groups (Sta, Sug, Fat, Pro, Fib) received 2.5% DSS in drinking water for 7 days. (A) Colon length in six groups. (B) The change of body weight in six groups on the seventh day. (C) DAI in six groups on the seventh day. (D) Normal control mice colon on day 7 showed well organized crypts and lamina propria and submucosal structures. (E), (F), (G), (H), (I): Histological analysis of representative colons from the mice in Sta, Sug, Fat, Fib and Pro groups, respectively. (original magnification, HE staining, 100 ×). *: P<0.05, compared to the Con group. #: P<0.05, compared to the Sta group. &: P<0.05, compared to the Fib group. a: P<0.05, compared to the Pro group.

**Fig 2.**
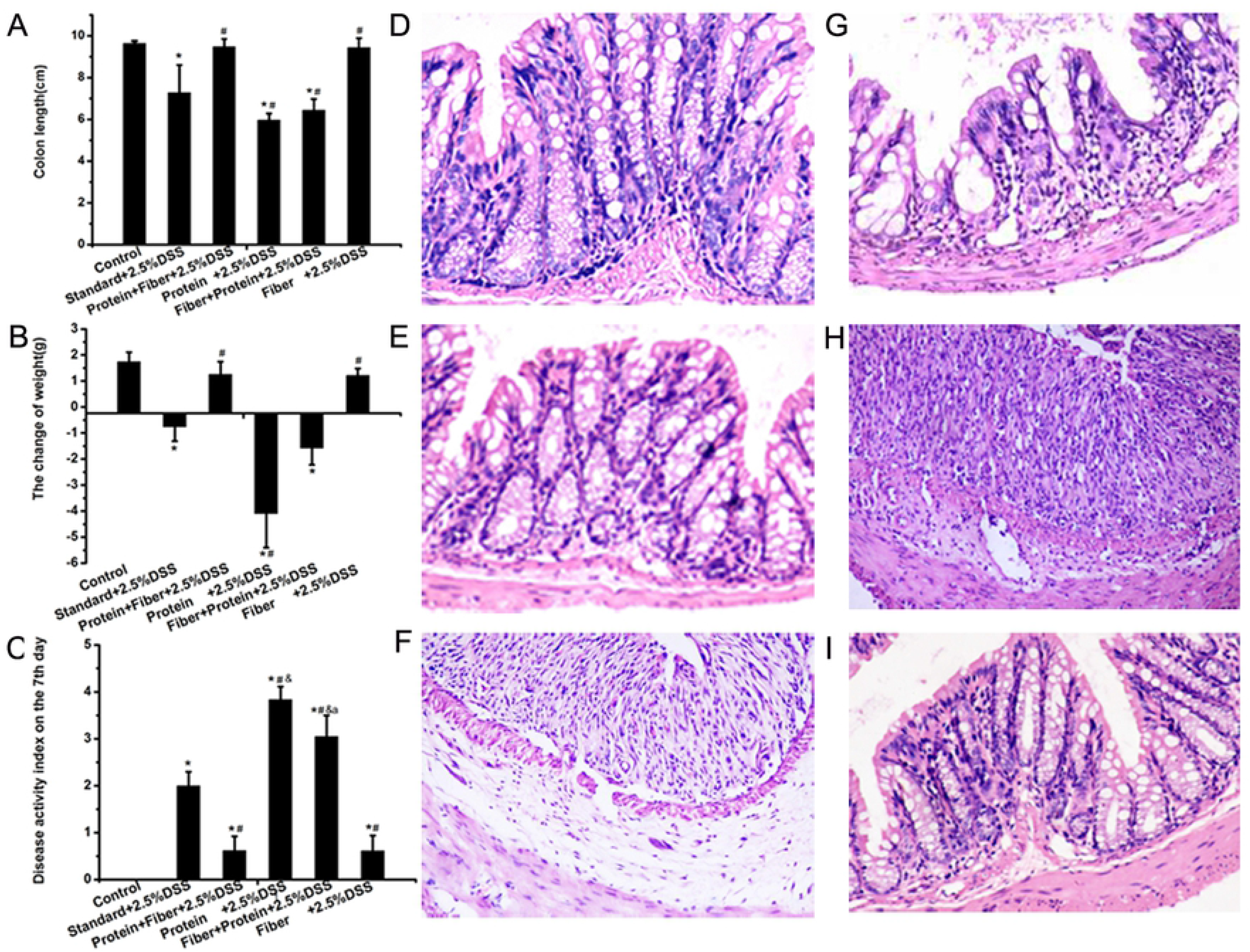
Effects of feeding order of high fiber and high protein on DSS induced colitis in the mice. Five groups (Sta, Pro+Fib, Pro, Fib+Pro, Fib) received 2.5% DSS in drinking water for 7 days. (A) Colon length in the six groups. (B) The change of body weight in all groups on the 7th day after drinking DSS. (C) DAI in six groups on the 7th day after drinking DSS. (D) Normal control mice colon on the 7th day showed well organized crypts and lamina propria and submucosal structures. (E), (F), (G), (H), (I), Histological analysis of representative colons from the mice in Sta, Pro+Fib, Pro, Fib+Pro and Fib groups, respectively. (original magnification, HE staining, 100 ×). *: P<0.05, compared to the Con group. #: P<0.05, compared to the Sta group. &: P<0.05, compared to the Fib group. a: P<0.05, compared to the Pro group.

**Fig 3.**
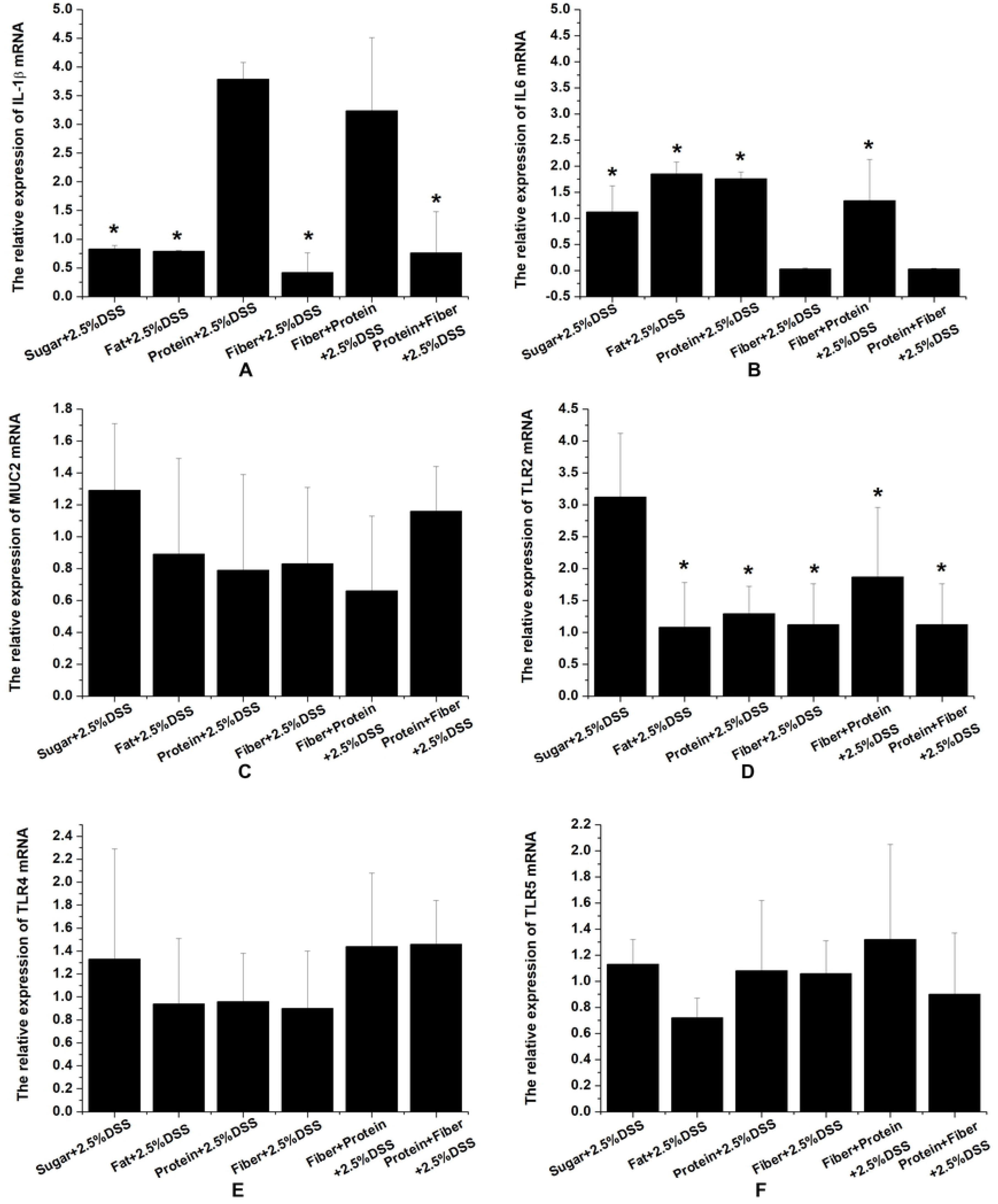
Effects of different diets on the expression of MUC2, TLR2, TLR4, TLR5, IL-1β and IL-6. A:The expression of IL-1βmRNA relative to the the Sta group after adding DSS, *: P<0.05, compared to the Pro group. B:The expression of IL-6 mRNA relative to the the Sta group after adding DSS, *: P<0.05, compared to the Fib group. C: The expression of MUC2 mRNA relative to the the Sta group before adding DSS. D: The expression of TLR2 mRNA relative to the the Sta group before adding DSS. *: P<0.05, compared to the Sug group. E: The expression of TLR4 mRNA relative to the the Sta group before adding DSS. F: The expression of TLR5 mRNA relative to the the Sta group before adding DSS.

### Microbial diversity in the gut of mice consuming different diets

To determine the changes of microbial diversity in mice fed with different diets we performed pyrosequencing on the stool of each group of mice. Pyrosequencing of 16S rRNA gene barcoded amplicons resulted in a total of 111,920 high-quality sequences, with a mean of 7,461 sequences (range 4,271-9,337) per sample. The Simpson index and ACE did not differ among different groups (P > 0.05). OTU (97%) of the Fat group was more than that of Pro and Fib group (P < 0.05), and OTU (97%) of the Fib group was less than that of the Sta group (Table 3, P < 0.05). Shannon index was significantly different in Fib group compared to the Sug and Fat group (Table 2, P < 0.05). Chao 1 was significantly different between Fib group and Fat group (Table 3, P < 0.05).

**Table 2.** OTUs (97%), alpha diversity (Shannon index, and Simpson index), richness (ACE and Chao1) for 16S rRNA libraries of stool.

| Group | OTU (97%) | Shannon<br>index | Simpson<br>index | ACE | Chao1 |
| --- | --- | --- | --- | --- | --- |
| Sta | 1175.67±279.14 | 4.79±1.26 | 0.08±0.10 | 3390.67±1049.89 | 2373.33±679.01 |
| Sug | 1120.33±220.51 | 5.38±0.50 | 0.02±0.01 | 3248.00±714.90 | 2268.33±426.16 |
| Fat | 1222.00±81.56 | 5.38±0.14 | 0.02±0.00 | 3356.00±304.87 | 2406.67±227.39 |
| Pro | 846.00±228.35 | 4.67±0.73 | 0.05±0.03 | 2548.33±993.02 | 1719.00±586.73 |
| Fib | 775.33±84.52 | 4.02±0.23 | 0.07±0.01 | 4144.67±1876.61 | 1551.67±95.87 |

From the Venn Diagram (Fig 4A), we can see that the OTU in the Pro group has 31.82% similarity with the Fib group, 31.72% similarity with the Fat group, 30.72% similarity with the Sug group, and 37.56% similarity with the other groups. The OTU in the Fib group has 43.35% similarity with the Pro group, and 46.28% similarity with the other groups. The OTU in the Fat group has 24.4% similarity with the Pro group, 37.39% similarity with the Sug group, and 51.64% similarity with the other groups. The OTU in the Sug group has 25.11% similarity with the Pro group, 39.73% similarity with the Fat group, and 51.39% similarity with the other groups.

**Fig 4.**
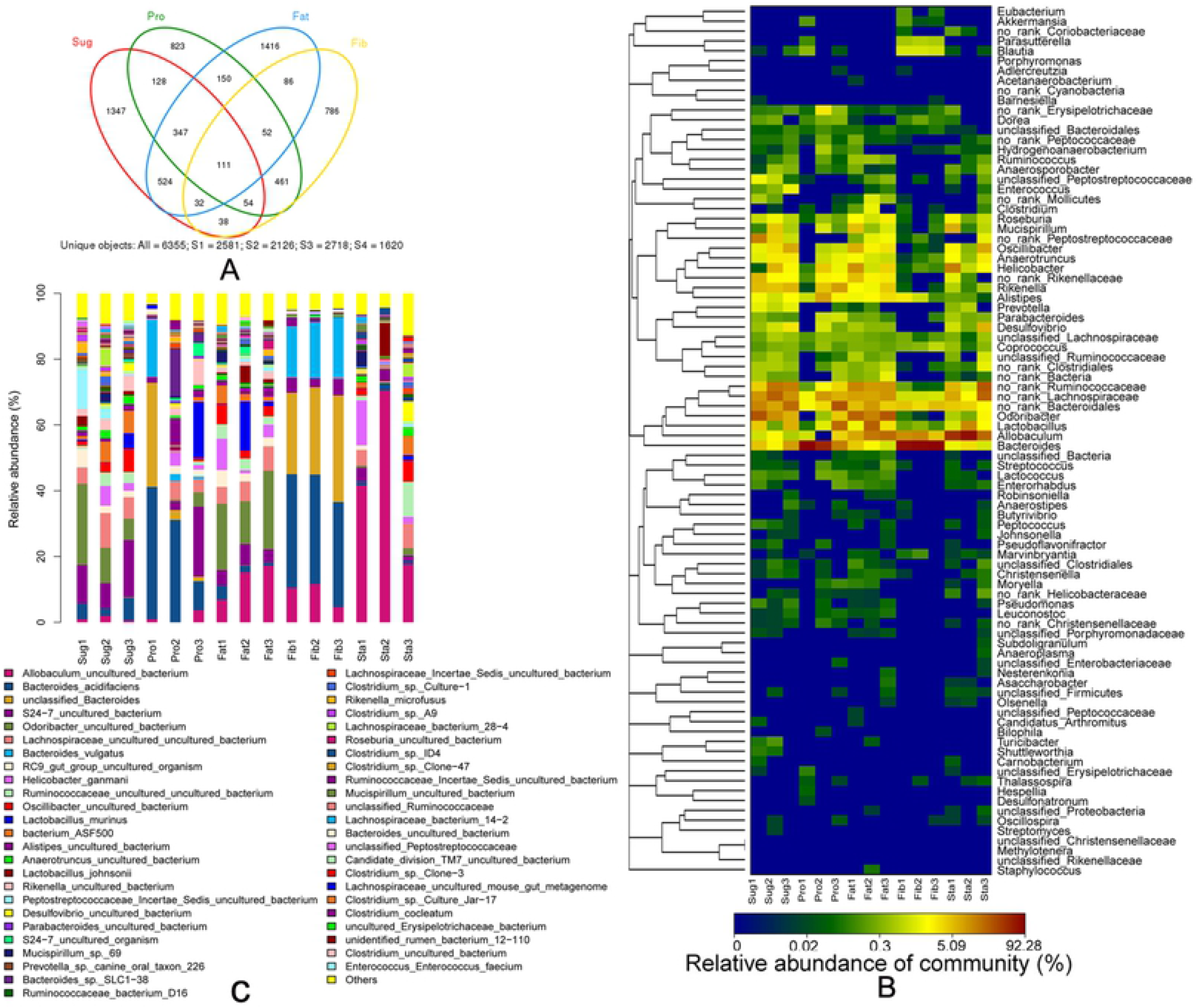
Microbial diversity in the gut of mice consuming different diets. Genomic DNA was extracted from the caecal content of randomly selected mice (3/group) and amplified using primers targeting the V1 to V3 hypervariable regions of bacterial 16S rRNA. (A) The Venn Diagram showed the differences and similarities in the Sug, Fat, Pro and Fib groups. (B)The heatmap showed the relative abundance of bacteria in the six groups (Con, Sta, Sug, Fat, Fib, Pro) in genus. Each column represented the abundance profile for a sample. (C) Bar chart showed the relative abundance of bacteria in the six groups (Con, Sta, Sug, Fat, Fib, Pro) in dominant species.

Different nutrients significantly affected the relative abundance of bacteria. *Bacteroidetes* and *Firmicutes* were the most abundant bacteria. Other predominant bacteria were *Actinobacteria, Deferribacteres, Proteobacteria, Tenericutes* and *Verrucomicrobia*. The amount of *Bacteroidetes* was the most in the Pro and Fib groups, and the least in fat and sugar groups (P < 0.05). In contrast, the amount of *Firmicutes* was the least in the Pro and Fib groups, and the most in fat and sugar groups (P < 0.05).

For bacterial genus, *Bacteroides* was the most abundant in Fib and Pro group, with no significant difference between them (P > 0.05). *Blautia* and *Parasutterella* was the most in Fib group compared with the other groups (P < 0.05). The amounts of *Blautia, Lactococcus* and *Parasutterella* were only 1.76% and 0.34% in the Fib and Pro group, respectively (P < 0.05) (Fig 4B). The composition of bacteria was the most similar between Fib and Pro groups, among all groups. For bacterial species, *Allobaculum uncultured bacterium* and *Bacteroides vulgatus* were the most in Fib group compared with the other groups (P < 0.05). There were 22 different species of bacteria between Pro and Fib groups (Fig 4C, P < 0.05), such as *Bacteroides vulgatus, Allobaculum uncultured bacterium, Blautia uncultured bacterium, Parasutterella uncultured bacterium, Blautia producta, Allobaculum uncultured bacterium* and *butyrate-producing bacterium L2-10*.

### Feeding order of high fiber and high protein led to the changes in microbial diversity

To further investigate whether the feeding order of dietary fiber or protein results in microbiota shift, we collected the stool of the Pro+Fib and Fib+Pro groups for Pyrosequencing.

From the Venn Diagram (Fig 5A), we can see that the OTU in the Pro+Fib group has 69.58% similarity with the Fib+Pro group, 55.85% similarity with the Pro group, 19.19% similarity with the Fib group, and 20.44% similarity with the other groups. The OTU in the Fib+Pro group has 76.90% similarity with the Pro+Fib group, 59.83% similarity with the Pro group, 18.10% similarity with the Fib group, and 13.97% similarity with the other groups. The OTU in the Fib group has 63.73% similarity with the Pro+Fib group, 54.40% similarity with the Fib+Pro group, and 9.84% similarity with the other groups. The OTU in the Pro group has 75.37% similarity with the Pro+Fib group, 73.05% similarity with the Fib+Pro group, and 5.89% similarity with the other groups.

**Fig 5.**
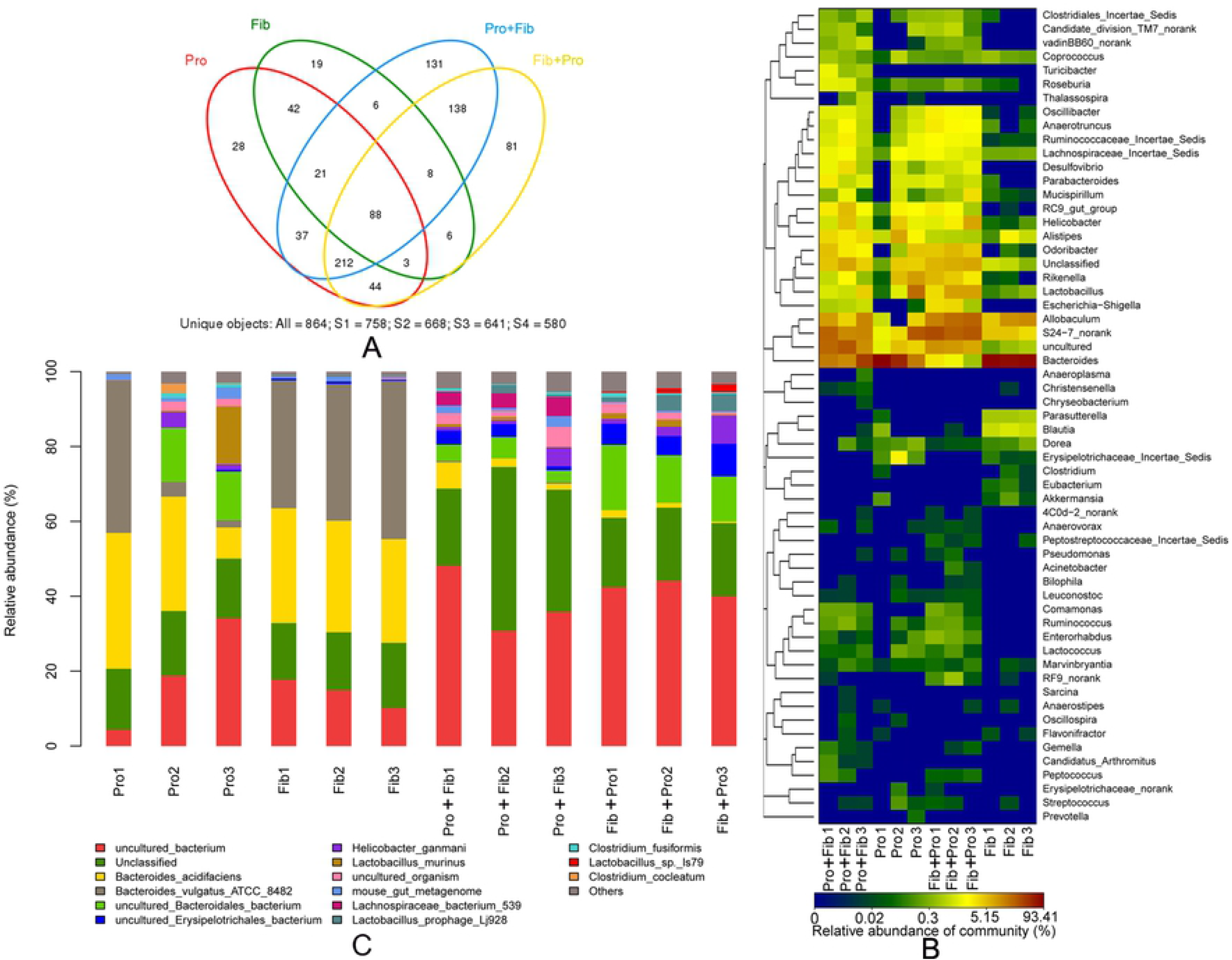
Microbial diversity after changing the feeding order of high fiber and high protein. Genomic DNA was extracted from the caecal content of randomly selected mice (3/group) and amplified using primers targeting the V1 to V3 hypervariable regions of bacterial 16S rRNA. (A) The Venn Diagram showed the differences and similarities in the Pro, Fib, Pro+Fib and Fib+Pro groups. (B) The heatmap showed the relative abundance of bacteria in the four groups (Pro+Fib, Pro, Fib+Pro, Fib) in genus. Each column represented the abundance profile for a sample. (C) Bar chart showed the relative abundance of bacteria in the four groups (Pro+Fib, Pro, Fib+Pro, Fib) in dominant species.

No significant difference in bacterial phylum was found between Pro+Fib and Fib+Pro groups. For bacterial genus, *Candidatus Arthromitus, Enterorhabdus, Escherichia-Shigella, RF9 norank* and *Turicibacter* were different in Pro+Fib and Fib+Pro group (Fig 5B, P < 0.05), with most significant change of *Escherichia-Shigella* and *Turicibacter*. In addition, 15 different species of bacteria were detected between Pro+Fib and Fib+Pro groups (Fig 5C, P < 0.05), such as *Lachnospiraceae bacterium 539, Lactobacillus sp. ls79* and *mouse gut metagenome*.

## Discussion

Living in sanitary environment and taking western-style diets at early life are considered to be risk factors for IBD, and important factors for shaping intestinal flora as well ^[25]^. Therefore, both dietary composition and gut microbiota play an important role in the pathogenesis of IBD. The aim of this study was to analyze that adult mice responsed to DSS and the structure of gut microbiota in newly weaned mice which fed with different components diets, in order to understand what dietary components and microbiota are associated with DSS-induced colitis. The results showed that mice fed with different dietary ingredients developed into three different disease phenotypes of DSS-induced colitis. The high protein diet exacerbates severity of DSS-induced acute colitis which characterized by weight loss, increased DAI scores, shortened colon length and increased histological scores. The high fiber diet had the effect of reducing colitis. The high sugar, high fat and standard diet showed the similar disease phenotype of colitis. These results suggested that mice fed different diets in early life respond differently to DSS in adult mice.

In previous studies, adult mice fed high sugar, high fat and Western-style diets have been found to aggravate DSS-induced colitis [26.2^7^]. However, in this study, adult mice fed with high sugar and high fat diets since early life did not aggravate DSS colitis. Obviously, the results were inconsistent between the two studies, and the difference between the two groups was the start time of high sugar and high fat diets feeding. It suggests that the colonic mucosa during the early stage of life is able to adapt with different diet. But, mutual adaptation does not explain the different responses of high protein and high fiber diets to DSS-induced colitis. In this study, weaning mice fed a high fiber diet for 4 weeks were converted to a high protein diet for 4 weeks, the result showed that the protective effect of the high fiber diet was lost and the high protein diet aggravated DSS-induced colitis. The other group which is converted the high protein diet to the high fiber diet without aggravating DSS-induced colitis, also showing only high fiber diet to reduce colitis. Previous studies found that eating high fiber diet for more than 20 days could enhance the protection of colonic mucosa ^[28]^ and this study shows that the time of giving 4 week a kind of the diet can counteract the effect of another early diet. Therefore, the high protein diet weakens or the high fiber diet strengthens resistance to DSS-induced colitis need to take time for developing the ability.

Natural immune receptors TLR2, TLR4, TLR5 and pro-inflammatory cytokines IL-6, IL-1 β in colon mucosa were detected. The levels of pro-inflammatory cytokines in the colon mucosa of mice were in the normal range before drinking DSS, and the levels of cytokines in colonic mucosa of DSS-exposed mice increased significantly, especially in the group which fed protein diet. These results suggest that early exposure to high protein and other unbalanced diets does not directly damage the mucosa. Some studies have shown that bacterial fermentation products of undigested proteins in colon are harmful to the mucosa ^[29–31]^. However, colitis was only aggravated when DSS is encountered in our study, which indicated the mechanism of bacterial fermentation is not suitable for mice which have adapted to high protein diet at the early stage of life.

The early stage of life is the critical period for the intestinal flora to mature into a stable adult microbiota. This study found that mice fed different diets after weaning had different phenotypes of DSS-induced colitis. Based on the clue of the disease phenotype, we tried to find out the relevant gut microbiota of DSS-induced colitis. Results showed that the gut microbiota of high fiber and high protein diets groups were similar, which is characterized by the decreased diversity and richness of the flora. The structure of gut microbiota had shown an interesting phenomenon that all mice which took the unbalance diets of high protein, fiber, fat, and sugar had very similar microbiota of the decreased abundance of *Firmicutes* and the increased abundance of *Bacteroides*. The microbiota alternation is similar to the dysbiosis in IBD patients, which is seems to be related to pathogenesis of IBD ^[32]^. In fact, the researches in both IBD patients and animal studies have shown that high fiber diet protect colon mucosa, and high protein diet aggravate colon mucosa damage ^[26,28,33]^. Obviously, the bacteria shaped by high fiber and protein diets in early life were not associated with IBD and DSS-induced colitis significantly. Because the evidence of dysbiosis in IBD mostly comes from the patients, and the structure of the microbiota cannot be known before the onset of the disease. In species/genus level, the result showed that the abundance of *Bacteroides vulgatus, Akkermansia uncultured bacterium* and *butyrate-producing bacterium L2-10* were higher in the mice with high fiber diet, these bacteria played beneficial roles in the colon^[34]^. This may partly explain why fiber supplement can decrease the severity of DSS induced colitis. On the other hand, high protein diets have been shown to increase the abundance of bacterial species that belonged to *Fusobacteriales, Coriobacteriales* and Clostridiales^[35]^. *Fusobacteriales* are implicated in many different inflammatory processes, including colonic inflammation ^[36,37]^. However, in this study there were no significant difference in *Fusobacteriales* in each group, which cannot explain why protein diet can aggravate the colitis. Therefore, the early adaptation of bacteria to intestinal mucosa may not damage the colonic mucosa. In our study, although we found that the bacteria between the mice with high fiber and the mice with high protein were the most similar, there were still some different bacteria between the two groups. Several of them, such as *Bacteroidesvulgatus, Akkermansia uncultured bacterium, butyrate-producing bacterium L2-10* and *Rikenella*, may play important roles in the development of colitis and further studies are needed to confirm their contributions to UC.

## Conclusion

Mice were fed with different dietary compositions of high sugar, fat, protein, and fiber since weaning, and similar intestinal flora of high-abundance *Bacteroides* and low-abundance *Firmicutes* are formed in adulthood. These microbiota do not cause colonic mucosal damage directly. Only a high protein diet aggravated DSS induced colitis, while high fiber diet alleviated the colitis.

